# Optimization of Energy State Transition Trajectory Supports the Development of Executive Function During Youth

**DOI:** 10.1101/424929

**Authors:** Zaixu Cui, Jennifer Stiso, Graham L. Baum, Jason Z. Kim, David R. Roalf, Richard F. Betzel, Shi Gu, Zhixin Lu, Cedric H. Xia, Rastko Ciric, Tyler M. Moore, Russell T. Shinohara, Kosha Ruparel, Christos Davatzikos, Fabio Pasqualetti, Raquel E. Gur, Ruben C. Gur, Danielle S. Bassett, Theodore D. Satterthwaite

## Abstract

Executive function develops rapidly during adolescence, and failures of executive function are associated with both risk-taking behaviors and psychopathology. However, it remains relatively unknown how structural brain networks mature during this critical period to facilitate energetically demanding transitions to activate the frontoparietal system, which is critical for executive function. In a sample of 946 human youths (ages 8-23 yr) who completed diffusion imaging as part of the Philadelphia Neurodevelopment Cohort, we capitalized upon recent advances in network control theory in order to calculate the control energy necessary to activate the frontoparietal system given the existing structural network topology. We found that the control energy required to activate the frontoparietal system declined with development. Moreover, we found that this control energy pattern contains sufficient information to make accurate predictions about individuals’ brain maturity. Finally, the control energy costs of the cingulate cortex were negatively correlated with executive performance, and partially mediated the development of executive performance with age. These results could not be explained by changes in general network control properties or in network modularity. Taken together, our results reveal a mechanism by which structural networks develop during adolescence to facilitate the instantiation of activation states necessary for executive function.

**SIGNIFICANCE STATEMENT:** Executive function undergoes protracted development during youth, but it is unknown how structural brain networks mature to facilitate the activation of the frontoparietal cortex that is critical for executive processes. Here, we leverage recent advances in network control theory to establish that structural brain networks evolve in adolescence to lower the energetic cost of activating the frontoparietal system. Our results suggest a new mechanistic framework for understanding how brain network maturation supports cognition, with clear implications for disorders marked by executive dysfunction, such as ADHD and psychosis.

## INTRODUCTION

Executive function is essential for a wide range of cognitive tasks, and is strongly associated with both overall intelligence (1) and academic performance (2). Executive function undergoes protracted maturation during adolescence (3, 4), and its development is linked to the expansion of the cognitive and behavioral repertoire. Notably, executive deficits are linked to both increased morbidity associated with risk-taking behaviors as well as a wide range of neuropsychiatric disorders (5), such as attention deficit hyperactivity disorder (ADHD) and psychosis (6, 7).

Prior studies have consistently established that executive function relies on activity in a distributed network of frontoparietal regions, including the dorsolateral prefrontal cortex, cingulate cortex, superior parietal cortex, and frontopolar cortex (8–12). Notably, both functional (13–16) and structural (17, 18) connectivity among these regions undergoes active remodeling during adolescence, with increased connectivity among executive regions, and diminished connectivity between executive regions and other systems such as the default mode network. As structural white matter networks are known to constrain both intrinsic connectivity and patterns of task-related activation (19, 20), it is possible that white matter networks develop during adolescence to facilitate dynamic transitions to activate the frontoparietal system. However, research that seeks to relate developing white matter networks to functional dynamics of executive function remains sparse.

Network control theory provides a powerful framework to address this gap in our knowledge. Specifically, network control theory allows one to integrate information regarding network topology and patterns of brain activation within one mathematical model, in order to specify how theoretical neural dynamics are constrained by the structural connectome (21). Such models assume that the activation state of the brain at a given time is a linear function of the previous state, the underlying white matter network, and any additional control energy injected into the system (22). From this paradigm, one can identify regionally specific control points that are optimally situated within the network’s topology to move the brain from one state to another (23–25). Network control theory thus could provide an account of how brain network topology facilitates patterns of brain activation. Previously, we have shown that the brain becomes more theoretically controllable during adolescence (18). While pivotal, that study considered only general properties regarding the controllability of a brain network. Innovations in network control theory (23, 24, 26) now allow one to describe how networks support transitions to specific activation states, including those required for executive function. We hypothesized that maturation of structural brain networks would allow for the target activation state of the frontoparietal executive system to be reached at a lower energetic cost.

To test this hypothesis, we capitalized on a large sample of youth who completed neuroimaging as part of the Philadelphia Neurodevelopmental Cohort (PNC) (27). We examined how white matter networks (estimated using diffusion imaging) support the transition to a frontoparietal system activation state. As described below, we demonstrate that the energy required to reach this state declines with age, especially within the frontoparietal control network. Furthermore, the whole-brain control energy pattern contains sufficient information to make accurate predictions about individuals’ brain maturity across development. Finally, participants with better performance on executive tasks require less energetic cost in the bilateral cingulate cortex to reach this activation target, and the energetic cost of this region mediates the development of executive performance with age. Notably, these results could not be explained by individual differences in network modularity or more general network control properties, and were not present in alternative activation target states. Together, these results suggest that during youth structural brain networks become optimized to minimize the energetic costs of transitions to activation states necessary for executive function.

## RESULTS

### Network topology constrains the transition to a frontoparietal activation state

In this study, we included 946 youths aged 8-23 years who were imaged as part of the PNC (**Fig. S1**). Structural white matter networks were reconstructed for each participant from diffusion imaging data using probabilistic tractography and a standard parcellation of 232 regions. Based on each participant’s unique network topology, we estimated the regional energetic cost required for the brain to transition from a baseline state to a frontoparietal activation target state (23, 24, 26) (**Fig. S2** & **Fig. 1A**). Formally, this estimation was operationalized as a multi-point network control optimization problem, where we aimed to identify the optimal trajectory between baseline and the frontoparietal activation target state that minimizes both the energetic cost and the distance between the final state and the target state.

**Fig. 1.**
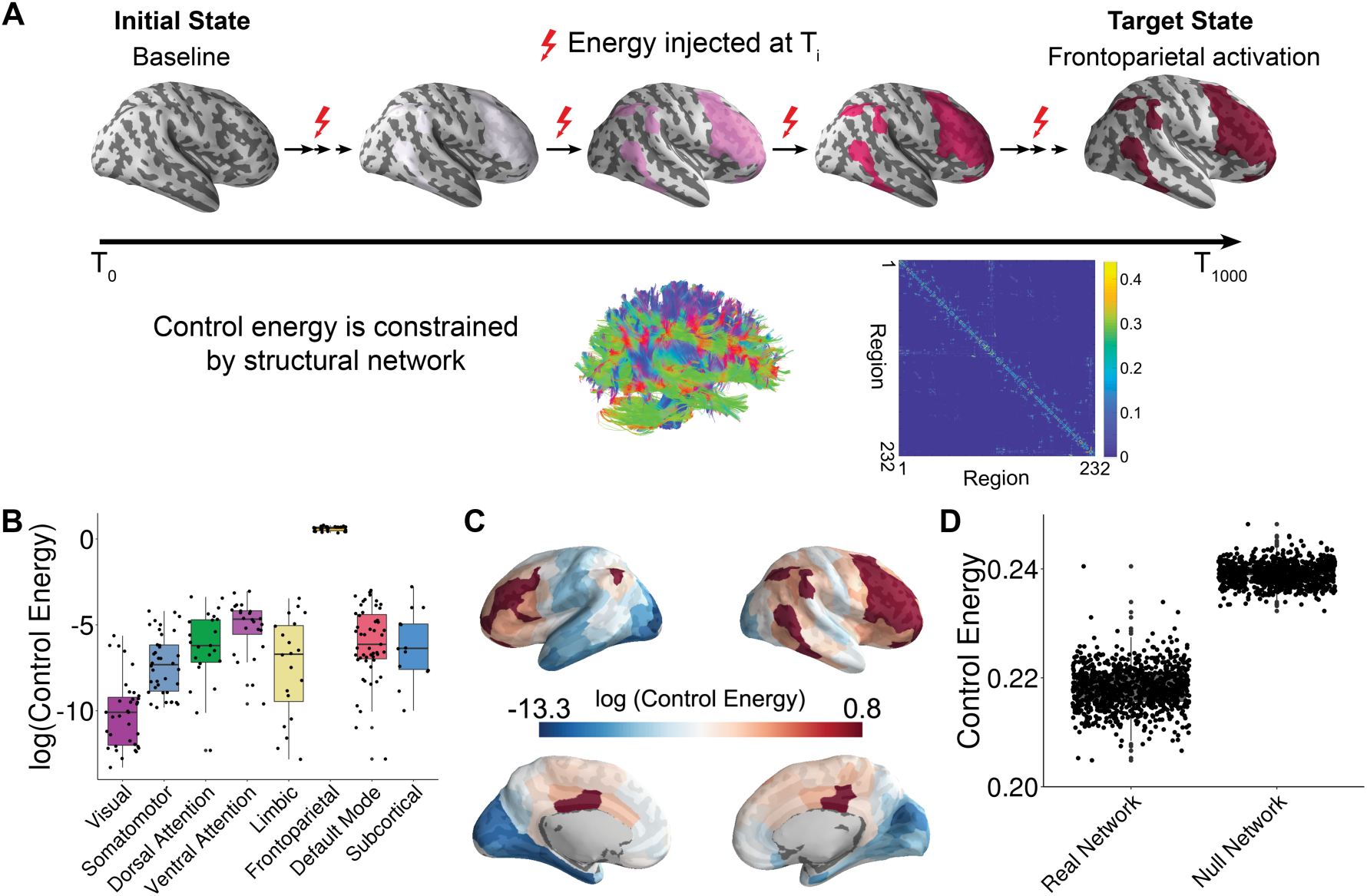
Schematic of the network control approach and the estimation of control energy. **A)** From a baseline state, we calculated the control energy required to reach a frontoparietal activation target state. This transition was calculated for each subject based on their structural brain network, which was estimated using diffusion imaging based probabilistic tractography. **B)** The average energetic costs to reach the frontoparietal activation target state varied by cognitive system, with the largest energetic costs being present in the frontoparietal control network and the ventral attention network. **C)** The regional control energy required to reach the frontoparietal activation target. **D)** The control energy cost of a transition to the frontoparietal activation target state was significantly lower in real brain networks than in null model networks where the strength and degree distribution were preserved.

Results of this dynamical systems model indicate that all participants arrived at the desired target state. For each network node and cognitive system, we calculated the mean control energy cost, which provides an indication of where energy must be injected into the network to achieve the transition to the target state. The highest control energy was observed in systems involved in executive function (**Fig. 1B** & **1C**), including the frontoparietal and ventral attention/cingulo-opercular systems (see **Fig. S2**) (28).

Based on recent evidence that network control properties depend appreciably on the topological structure of the network (29), we next sought to demonstrate that the topological structure of brain networks specifically facilitates this transition. We therefore compared the energetic cost of this transition in empirical brain networks to the energetic cost observed in null model networks. Specifically, we randomly permuted (100 times per participant) the placement of edge weights, while preserving the network degree and strength distribution. The mean whole brain energetic cost of the null networks was significantly higher (*t* = 180.08, *p* < 2 × 10^−16^) than that of the empirical networks (**Fig. 1D**), indicating that structural brain networks are topologically optimized to reduce the energetic costs of the transition to a frontoparietal activation state.

### Energetic costs of the transition to a frontoparietal activation state decline with development

Having shown that the topology of structural brain networks facilitates transitions to a frontoparietal activation state, we next investigated how the energetic costs of this transition evolve in youth. We hypothesized that the energy required to make this transition would decline as networks were remodeled in development. To test this hypothesis, we used generalized additive models (GAM) with penalized splines to examine both linear and nonlinear associations of control energy and age within a statistically rigorous framework. Age associations with control energy were examined at multiple scales, including the level of the whole brain, cognitive systems, and individual nodes. For all analyses, we included sex, handedness, in-scanner head motion, total brain volume, and total network strength as covariates. These analyses revealed that the whole-brain average energetic cost of the transition to the frontoparietal activation state declined with age (*P* = 3.06 × 10^−7^) (**Fig. 2A**). Notably, analyses of cognitive systems indicated that age effects were heterogeneously distributed (**Fig. 2B)**, with the largest declines in control energy occurring in frontoparietal (*P*_FDR_ = 4.54 × 10^−7^; **Fig. 2C**), visual (*P*_FDR_ = 5.71 × 10^−5^), and somatomotor (*P*_FDR_ = 2.70 × 10^−3^) systems. In contrast, energetic costs within the limbic (*P*_FDR_ < 2 × 10^−16^) and default mode (*P*_FDR_ = 5.66 × 10^−3^) systems increased with age (see **Fig. S3**). These system-level results aligned with analyses of individual network nodes; we found that the control energy of 49 regions decreased significantly with age (*P*_FDR_ < 0.05), including regions in the frontoparietal control network, visual network, and somatomotor network. Furthermore, the control energy significantly increased with development in 30 regions (*P*_FDR_ < 0.05), which were mainly situated in limbic and default mode systems (**Fig. 2D**).

**Fig. 2.**
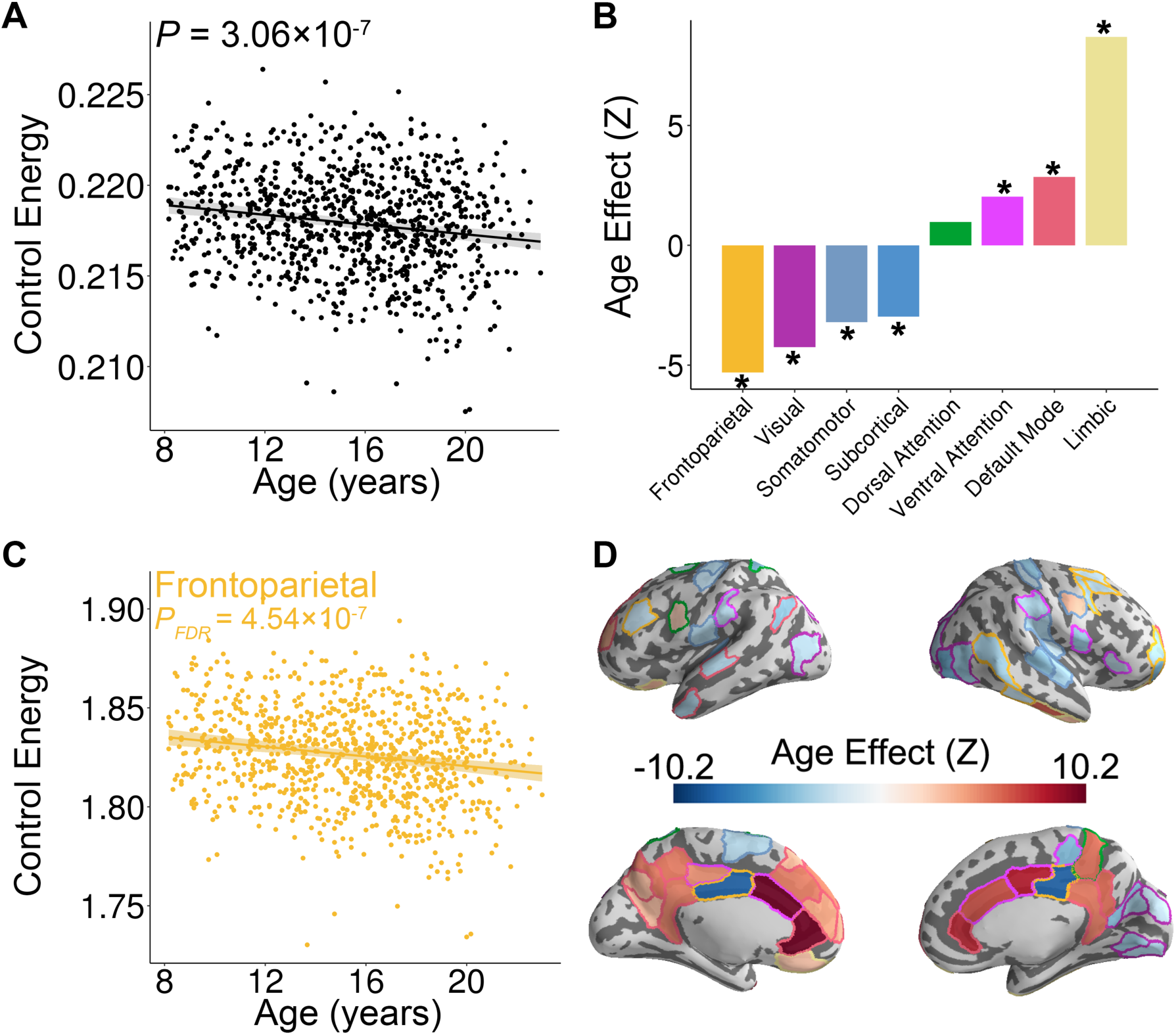
Control energy evolves with age in youth. **A)** The mean whole-brain control energy required to reach the frontoparietal activation target declined with age. **B)** Control energy declines significantly with age in the frontoparietal, visual, somatomotor and subcortical systems. In contrast, control energy increased in the ventral attention, default mode and limbic systems. **C)** The control energy of the frontoparietal system declines significantly with age. **D)** The age effect of control energy for each brain region. The color of the contour of each brain region represents the cognitive system for each region (see **Fig. S2**). In the scatterplots shown in panels **A** and **C**, data points represent each subject (n = 946), the bold line indicates the best fit from a general additive model, and the shaded envelope denotes the 95% confidence interval.

### Specificity and sensitivity analyses provide convergent results

Having found strong associations between age and control energy, we next conducted a series of four additional analyses to evaluate whether these results could be attributed to other network properties. First, we evaluated whether age effects could be due to non-topological network properties by evaluating the presence of age effects in null networks where degree and strength distributions were preserved. We found that the significance level of age effects in null networks were smaller than those observed in the real network (*p* < 0.01, 100 permutations), suggesting that the empirically measured developmental effects were indeed driven by changes in the network topology (**Fig. S4A**). Second, we determined whether these developmental effects were only associated with specific activation target states. Notably, declines in control energy with age were more significant (*p* = 0.03, 100 permutations) than those observed when randomized activation target states were used (**Fig. S4B**).

Third, we evaluated the specificity of these developmental effects by evaluating age effects in the transition to an *a priori* motor system activation target (**Fig. S2**) (28). As the age range of 8-23 years is a critical period in the development of executive function rather than motor function, we expected weaker age effects in the motor system. We found that the whole-brain control energy required to transition to the motor system activation did not significantly change over the age range studied (*P* = 0.14, **Fig. S4C**). Fourth, we evaluated whether our developmental results could be explained by other network properties known to change with development, including modal controllability (18) or network modularity (17). Notably, including these properties as model covariates did not alter our results (**Fig. S5**). For example, average control energy of the whole-brain and frontoparietal system both significantly declined with age after controlling for modal controllability (whole-brain: *P* = 2.09 × 10^−9^; frontoparietal: *P*_FDR_ < 2 × 10^−16^) as well as after controlling for network modularity (whole-brain: *P* = 7.73 × 10^−5^; frontoparietal: *P*_FDR_ = 6.46 × 10^−5^).

### Patterns of control energy can predict brain maturity

Having established that control energy required to reach the frontoparietal activation state changes with age on a regional and system-level basis, we asked whether the multivariate pattern of control energy could be used to identify an individual participant’s age in an unbiased fashion. To address this question, we used ridge regression with nested two-fold cross validation (2F-CV, see **Fig. S6**). Specifically, we divided all subjects into two subsets based on age (30, 31), with the first subset used as a training set and the second subset used as a testing set. Within the training set, we used inner 2F-CV to select an optimal regularization parameter (*λ*). Then, we trained a model using the training data and predicted the brain maturity (i.e., ‘brain age’) of participants in the testing set (32, 33). The significance of the model was evaluated using permutation testing, where the correspondence between a subject’s brain network and their age was permuted uniformly at random. This analysis revealed that the multivariate pattern of control energy could accurately predict an unseen individual’s age with a high degree of accuracy (**Fig. 3A** and **Fig. S7A** & **S7B**): the correlation between the predicted ‘brain age’ and chronological age was 0.62 (*p* < 0.001), while the mean absolute error (MAE) was 2.27 years (*p* < 0.001). For completeness, we also repeated this procedure while reversing the training and test sets, which yielded very similar results (*r* = 0.57, *p* < 0.001; MAE = 2.28, *p* < 0.001) (**Fig. 3A** and **Fig. S7C** & **S7D**). When model weights were examined at the level of individual network nodes, the regions that most contributed to the prediction of brain maturity aligned with univariate analyses, and included the dorsolateral and ventrolateral prefrontal cortex, the cingulate cortex, superior parietal cortex, and lateral temporal cortex (**Fig. 3B**). In order to ensure that our initial split of the data was representative, we repeated this analysis with 100 random splits, which returned highly consistent results (mean *r* = 0.59, mean MAE = 2.27 years).

**Fig. 3.**
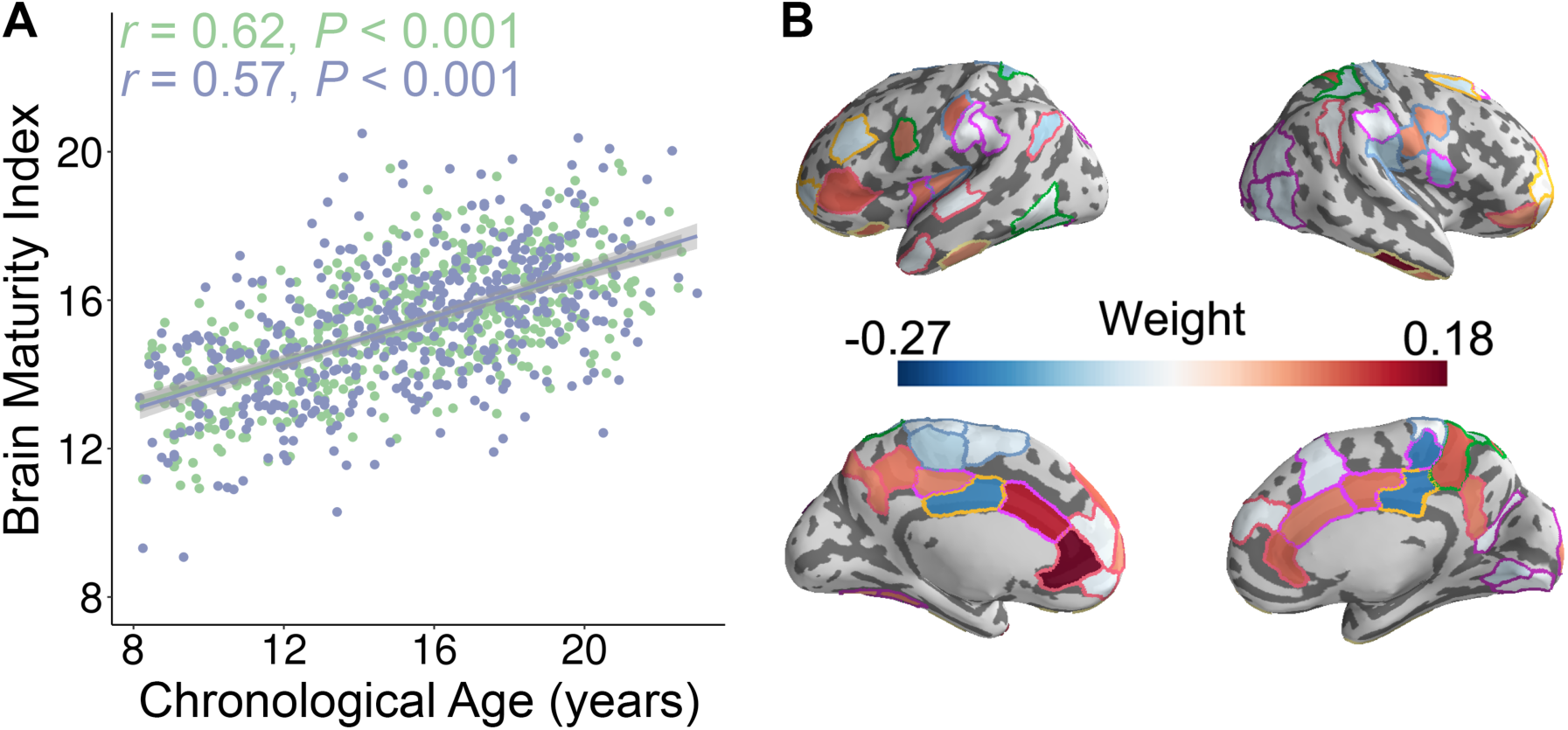
The whole-brain control energy pattern contains sufficient information to accurately predict brain maturity in unseen individuals. **A)** The predicted brain maturity index was significantly related to the chronological age in a multivariate ridge regression model that used 2-fold cross validation (2F-CV) with nested parameter tuning. The group of all subjects was divided into two subsets according to age rank; all models were trained using a completely independent sample of the data. The blue color represents the best-fit line between the actual score of the first subset of subjects and their scores predicted by the model trained using the second subset of subjects. The green color represents the best-fit line between the actual score of the second subset of subjects and their scores predicted by the model trained using the first subset of subjects. **B)** Regions with the highest contribution to the multivariate model aligned with mass-univariate analyses and included frontal, parietal, and temporal regions.

### Declines in control energy mediate the development of executive function

Lastly, we investigated the cognitive implications of individual differences in control energy. Specifically, we expected that participants with higher executive performance on a standardized cognitive battery would require reduced control energy cost to activate the frontoparietal system. In order to ensure that associations were present above and beyond the observed developmental effects, we controlled for linear and nonlinear age effects in addition to the other covariates described above. While we did not find effects at the whole-brain or systems level, we did find that reduced control energy within two regions in the frontoparietal control system – the left and right middle cingulate cortex – was associated with higher executive function (Left: *P_FDR =_* 0.032; Right: *P_FDR =_* 0.002. **Fig. 4A & 4B**).

**Fig. 4.**
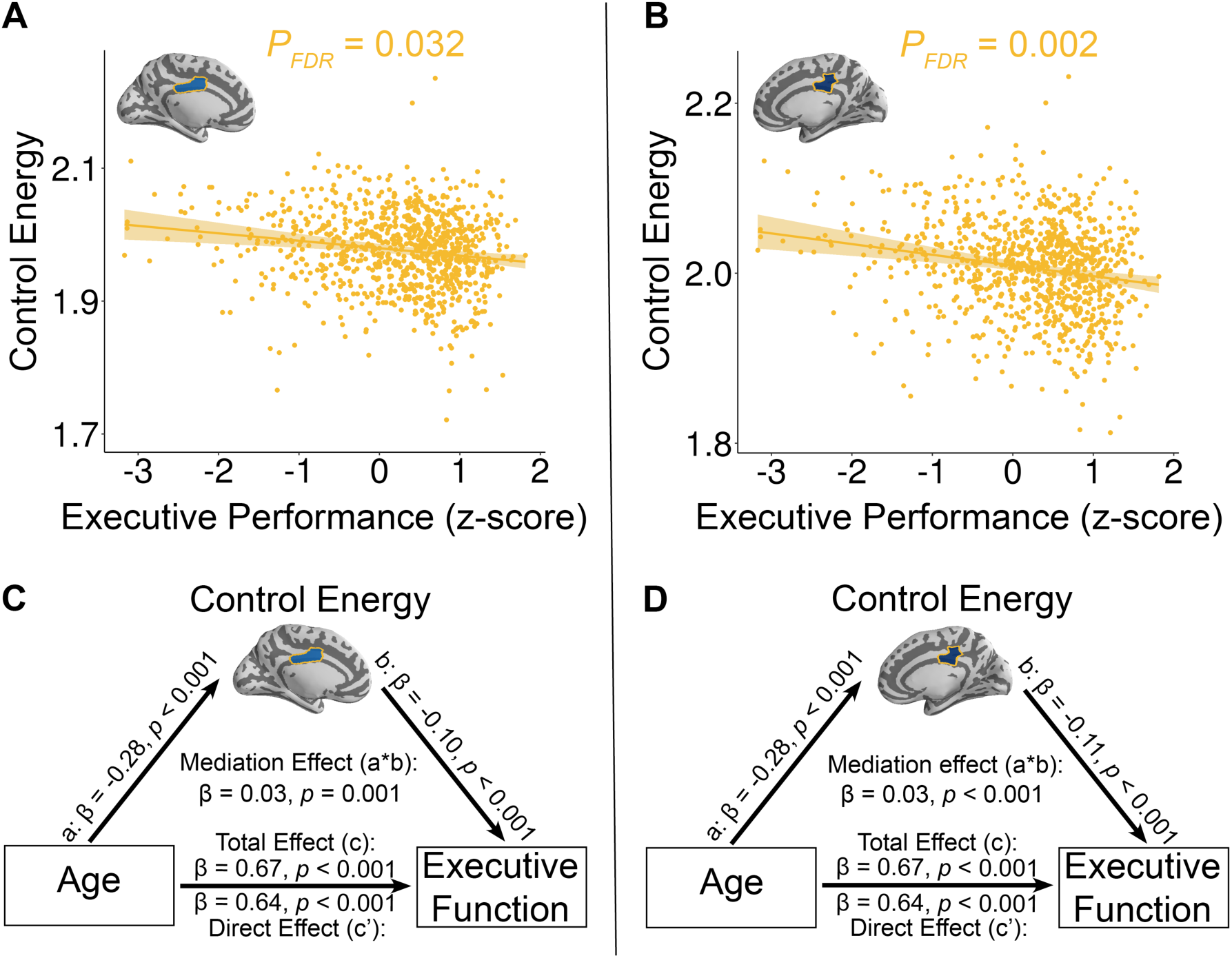
Reduced control energy in both the **A)** left and **B)** right mid-cingulate cortex is associated with higher executive performance. Data points represent each subject (n =944), the bold line indicates the best linear fit, and the shaded envelope denotes the 95% confidence interval. The yellow color indicates that the two regions belong to the frontoparietal system (**Fig. S2**). The control energy of both **C)** left and **D)** right mid-cingulate cortex partially mediates the improvement of executive function with age. Significance of mediation effect was assessed using bootstrapped confidence intervals.

Having identified a significant relationship between control energy in the bilateral middle cingulate cortex and both age and executive function, we conducted mediation analyses to investigate the extent to which control energy accounted for the association between age and executive function. Using a bootstrapped mediation analysis while adjusting for the covariates described above, we found that control energy in both the left (β = 0.03, *p* = 0.001, 95% confidence interval = [0.01, 0.04]; **Fig. 4C**) and right middle cingulate cortex (β = 0.03, *p* < 0.001, 95% confidence interval = [0.02, 0.05]; **Fig. 4D**) mediated the development of executive function with age.

## DISCUSSION

Using a large sample of youths and a mechanistic model of brain network function, we demonstrated that the control energy theoretically required to transition to a frontoparietal activation state declines with age in youth. Furthermore, the multivariate pattern of the whole-brain control energy predicted the brain maturity of unseen individual participants. These results could not be explained by individual differences in network modularity or other network control properties, and were not observed in analyses stipulating alternative activation targets. Finally, we found that individuals who had higher executive function required lower control energy in the bilateral middle cingulate cortex to activate the frontoparietal system, and the control energy of this region partially mediated the development of executive performance with age. These results suggest that maturation of structural brain networks facilitates the emergence of executive dynamics in youth.

In this study, we used a model of multi-point control to understand the transition from a baseline state to a target state where the frontoparietal system is activated. Building upon prior work (23–26), this dynamical system was constrained by the brain’s structural connectome. Across all participants (regardless of age), we found that the energetic cost of this transition was far lower in real brain networks compared to null networks that preserved basic properties such as degree and strength. This result extends prior work demonstrating that the topology of a network has marked implications for its control properties (25, 29). In the context of human cognition, the results suggest that brain network topology is configured to support these energetically demanding transitions to activation states required for executive function.

We evaluated associations between control energy and age at multiple spatial and topological scales, including the whole brain, cognitive systems, and individual network nodes. We observed that the mean whole-brain control energy required to transition to the frontoparietal activation state declines with development. This observation suggests that white matter network topology is optimized during youth to facilitate energetically efficient transitions to activation states that are necessary for executive function.

Examination of individual cognitive systems revealed that this decline in whole-brain energy was driven by reduced energetic costs within the frontoparietal system. In particular, substantial negative associations between age and control energy were observed in lateral prefrontal cortex and middle cingulate cortex. These regions are responsible for preparation, execution, monitoring and switching of tasks (i.e., working memory, attention, inhibitory control, etc.) (8, 9, 11, 34). The reduced regional energetic cost suggests that structural brain networks may mature to allow for neural events that occur in these regions to impact the broad activation state of the entire network more efficiently, and more easily drive the brain towards the frontoparietal activation state associated with demanding executive tasks (15, 17, 35).

In contrast, the energetic cost of limbic regions and regions within the default mode system increased with age. This localization of costs suggests that these regions become less able to move the brain to a frontoparietal activation state as development progresses. This result is consistent with previous studies using both structural and functional connectivity data, which have shown that the frontoparietal system becomes more segregated from the default mode during youth (14, 17), potentially allowing for functional specialization and reducing interference. However, developmental changes in modularity do not explain our current findings, as sensitivity analyses that included modularity quality as a covariate yielded convergent results.

It should be noted that in prior work we demonstrated that another network control property – modal controllability – increased with age (18). That previous study used a single-point model of control, where modal control quantified the ability of a single brain region to move the brain to any theoretically possible state (22). Here, we build upon those prior results and show that in the context of multi-point control, energetic costs to a specific frontoparietal activation target state decline with age. By allowing for multi-point control and stipulating a specific activation target, these results are both consistent with our prior result and also more biologically interpretable. Importantly, we demonstrate that our current results are not simply a result of increasing single-point modal controllability: when modal controllability was included as a model covariate, our results remained unchanged. Thus, our results cannot be easily explained by general control properties of the network that are known to evolve in development.

Beyond single-point modal control and modularity, we also evaluated whether other confounds or network properties could explain our developmental findings. Specifically, we found that null networks that preserved other basic network properties (i.e., degree and strength) did not show similar age effects. Analogously, we observed much stronger declines in network control energy for the frontoparietal activation target than when randomly shuffled activation targets were specified. However, while such shuffled activation targets provide a useful null condition, they have limited biologic plausibility. Accordingly, we also evaluated age effects using a somatomotor activation target constructed using the same atlas as our frontoparietal target. Notably, no developmental associations with control energy were seen with this somatomotor target. These results accord with prior evidence indicating that motor development precedes executive development, and is largely complete by late childhood or early adolescence (36). Moreover, these results suggest that the observed developmental changes in control energy may be specific for transitions to activation states recruiting higher-order cognitive systems, which undergo protracted maturation.

Our main results regarding brain development (as well as the supplementary analyses described above) used a mass-univariate analysis approach, where the association between the control energy of each region was modeled separately. Complementary analyses sought to identify distributed multivariate patterns, which could be used to predict the brain maturity of unseen individuals. Such an approach is similar to prior studies that have used structural (33, 37), functional (16, 32), or diffusion (38) based imaging to predict brain development. Here, we used a rigorous split half validation framework with nested parameter tuning. We found that the complex pattern of control energy could be used to predict individual brain maturity with a relatively high degree of accuracy. The feature weights from this multivariate model were generally consistent with findings from mass-univariate analyses, underscoring the robustness of these results to the methodological approach. With the ability to predict individuals’ brain maturity, control energy could have potential to determine whether individuals display either precocity or delay in specific dynamic aspects of brain maturation, which may be relevant to studying developmental disorders and neuropsychiatric syndromes (32, 38).

Having established that control energy evolves significantly in youth, as a final step we evaluated whether control energy has implications for individual differences in executive performance. While controlling for age, we observed a significant negative correlation between control energy of both the bilateral middle cingulate cortex and executive function performance. The middle cingulate is a component of the frontoparietal control system (15, 28) and is critical for executive tasks such as performance monitoring, error detection, and task switching (34, 39). This result suggests that individuals with better executive function can transition to the frontoparietal activation state while making fewer demands on this region. Critically, the control energy of the bilateral middle cingulate cortex mediated the observed improvement of executive function with age.

Several limitations should be noted. First, all data presented here were cross-sectional, which precludes inference regarding within-individual developmental effects. Ongoing follow-up of the PNC will yield informative longitudinal data, as will other large-scale studies such as the Adolescent Brain and Cognitive Development Study. Second, it should be noted that probabilistic tractography methods remain limited in their ability to fully resolve the complex white matter architecture of the human brain. However, these methods are currently considered state-of-the-art, and may be superior to tensor-based tractography in resolving crossing fibers (40). Third, we used a linear model of brain network dynamics, which has been shown to be an appropriate approximation in past efforts (22, 25, 41). However, future studies could expand upon this work and use more complex non-linear models (42). Finally, it should be noted that motion artifact is a major potential confound for any study of brain development, and prior studies by our group and others have shown that motion artifact can bias estimates of tractography and confound developmental inference (43). However, to limit the impact of this confound, we conducted rigorous quality assurance and included in-scanner motion as a covariate in all analyses.

These potential limitations notwithstanding, we demonstrated that the topological structure of white matter networks is optimized during development to facilitate transitions to a frontoparietal activation state. Moving forward, this framework may be useful for understanding the developmental substrates of executive dysfunction in diverse psychiatric disorders including psychosis and ADHD. Improved knowledge regarding both normal network development and abnormalities associated with psychopathology is a prerequisite for developing individualized interventions to alter disease trajectories and improve patient outcomes. In the future, advances in non-invasive neuromodulatory therapies may allow for targeted stimulation of specific brain regions that are optimally situated within the brain’s control architecture to facilitate transitions to specific target states (44). Such advances could potentially aid in the treatment of the wide range of neuropsychiatric disorders marked by executive dysfunction (45).

## MATERIALS AND METHODS

### Participants and Assessments

All subjects or their parent/guardian provided informed consent, and minors provided assent. The Institutional Review Boards of both Penn and CHOP approved study procedures. Following exclusion of participants with major medical disorders, missing data, or low-quality imaging data (see SI Methods), we included a sample of 946 subjects aged 8-23 years (27).

### Cognitive Assessment

All participants completed the Penn Computerized Neurocognitive Battery (3). Accuracy and speed measures were summarized using factor analysis to yield a performance efficiency score for executive function (46); see SI Methods for details.

### Image Acquisition and Preprocessing

All participants completed multi-modal neuroimaging on the same scanner using the same sequences (27). Diffusion images were pre-processed using standard procedures, and probabilistic tractography (40, 47) was used to construct a structural connectivity matrix among 232 regions defined using a standard surface-based structural parcellation (48–50). We treat this connectivity matrix as a parsimonious encoding of a network model (51). Edge weights within this network were defined as the connection probability between any pair of nodes (43, 52, 53). In contrast to tensor-based deterministic approaches, probabilistic tractography allowed us to model up to two crossing fibers per voxel, enhancing sensitivity to more complex white matter architecture (40). As described previously (17), each network node was assigned to an *a priori* network community using a commonly-used functional atlas (28). Details are available in SI Methods.

### Calculation of control energy

We calculated control energy using previously described methods (23, 24, 26) and standard parameters (see SI Methods). Specifically, based on the structural connectivity network of each subject, we evaluated the control energy of all brain regions necessary to move each from a baseline state to a target state with activation of the frontoparietal system. The control set included all regions in the parcellation.

### Comparison to null model network

In order to determine whether the topology of brain networks specifically facilitated transitions to the frontoparietal activation target state, we compared the energetic cost to that of null model networks. Specifically, for each participant we constructed 100 null model networks where the degree and strength distribution was preserved (54). We compared the control energy cost of the transition to the frontoparietal activation target state estimated from the empirical networks to the average energy cost estimated in these null networks using a paired *t*-test.

### Statistical analyses of developmental and cognition effects

Brain development is known to be a nonlinear process. Accordingly, for our developmental analyses we used generalized additive models (GAMs) in order to simultaneously model linear and nonlinear relationships with age using penalized splines (55). We evaluated associations between control energy and age at multiple resolutions, including the whole brain, cognitive systems, and network nodes. Similarly, we evaluated associations between control energy and executive performance while controlling for age. Furthermore, for regions that displayed both cognition and age associations, we evaluated whether regional control energy might mediate the relationship between age and executive function using a boostrapped mediation analysis (resampled 10,000 times). For all models, we included sex, handedness, total brain volume, total network strength, and in-scanner head motion during the diffusion scan as model covariates. Multiple comparisons were accounted for using the False Discovery Rate (*Q*<0.05).

### Individualized prediction of brain maturity

As a complement to the mass-univariate analyses described above, we also sought to predict individual brain maturity using the multivariate pattern of control energy (32, 33, 38). We used ridge regression (31) with nested two-fold cross validation (2F-CV). In this model, the outer 2F-CV was used for testing and the inner 2F-CV was used for parameter selection (*λ*) (**Fig. S6**). Specifically, we divided the dataset into two subsets according to the age rank (30, 31), and we used one subset as the training set and the other subset as the testing set. We regressed out the covariates from each feature for the training set; these coefficients were then applied to the testing set. We selected an optimal parameter *λ* with an inner 2F-CV and then trained a model including all training subjects using the acquired optimal *λ* This model was used to predict individuals’ brain maturity (i.e., ‘brain age’) of the left-out testing set. This procedure was repeated for each half of the split data. The correlation and mean absolute error (MAE) between the predicted ‘brain age’ and chronological age was used to quantify the degree to which the model captured the development trajectory of the brain. To verify that the split of our data (based on age rank) was representative, we also repeated the analysis described above using 100 random splits.

### Specificity and sensitivity analysis

We conducted several additional supplementary analyses to assess the sensitivity and specificity of our results. First, to ensure the observed associations with age were driven by the topological structure of real brain networks, we tested whether age effects existed using null networks. Second, we tested whether the observed age effects were specific to the frontoparietal target state, or were also present using randomly shuffled target states. Third, we evaluated the developmental effects of control energy with a motor activation target state to evaluate the specificity of our observed age effects. Fourth, to ensure that observed associations were not simply dependent on potentially related network properties, we re-examined associations with age while considering modal controllability or overall network modularity as a covariate. See SI for details.

### Data & code availability

The PNC data is publicly available in the Database of Genotypes and Phenotypes: accession number: phs000607.v3.p2; https://www.ncbi.nlm.nih.gov/projects/gap/cgibin/study.cgi?study_id=phs000607.v3.p2. All analysis code is available here: https://github.com/ZaixuCui/pncControlEnergy, with detailed explanation in https://github.com/ZaixuCui/pncControlEnergy/wiki.

## ACKNOWLEDGEMENTS

Thanks to Chad Jackson for data management and systems support. This study was supported by grants from National Institute of Mental Health: R21MH106799 (D.S.B. & T.D.S.), R01MH107703 (T.D.S.), R01MH112847 (R.T.S. & T.D.S.), R01MH107235 (R.C. G.), R01MH113550 (T.D.S. & D.S.B.), and R01EB022573 (C.D.). The PNC was supported by MH089983 and MH089924. Additional support was provided by P50MH096891 to R.E.G., K01MH102609 to D.R.R., R01NS085211 to R.T.S., and the Dowshen Program for Neuroscience. Additionally, D.S.B. acknowledges support from the John D. and Catherine T. MacArthur Foundation, the Alfred P. Sloan Foundation, the ISI Foundation, the Paul Allen Foundation, the Army Research Laboratory (W911NF-10-2-0022), the Army Research Office (Bassett-W911NF-14-1-0679, Grafton-W911NF-16-1-0474, DCISTW911NF-17-2-0181), the Office of Naval Research, National Institute of Health (2-R01-DC-009209-11, 1R01HD086888-01, R01 – MH112847, R01-MH107235), National Institute of Neurological Disorders and Stroke (R01 NS099348), and National Science Foundation (BCS-1441502, BCS-1430087, NSF PHY-1554488 and BCS-1631550). The content is solely the responsibility of the authors and does not necessarily represent the official views of any of the funding agencies.

## Author Contributions

All authors contributed to this work, approved the final manuscript, and agreed to the copyright transfer policies of the journal.

## Supplementary Information

### MATERIALS AND METHODS

#### Participants

Overall, 1,601 participants were enrolled. However, 340 subjects were excluded owing to clinical factors including medical disorders that could affect brain function, current use of psychoactive medications, prior inpatient psychiatric treatment, or an incidentally encountered structural brain abnormality. Among the 1,261 subjects eligible for inclusion, 54 subjects were excluded for a low quality T1-weighted image or errors in the FreeSurfer reconstruction. Of the remaining 1,207 subjects with a usable T1 image, 128 subjects were excluded because of the lack of a complete diffusion scan. Of the 1,079 subjects with complete diffusion data, 110 subjects failed quality assurance as part a rigorous quality assurance protocol for diffusion MRI (1). Additionally, 20 subjects were excluded because they had no field map for distortion correction. Finally, of the remaining 949 subjects, 3 subjects were excluded due to incomplete image coverage during brain parcellation, yielding a final sample of 946 participants (**Fig. S1**).

#### Cognitive assessment

The Penn computerized neurocognitive battery (Penn CNB) was administered to all participants during a separate session from neuroimaging. The CNB consists of 14 tests adapted from tasks applied in functional neuroimaging to evaluate a broad range of cognitive domains (2). These domains include executive control (abstraction and mental flexibility, attention, working memory), episodic memory (verbal, facial, spatial), complex cognition (verbal reasoning, nonverbal reasoning, spatial processing), social cognition (emotion identification, emotion differentiation, age differentiation) and sensorimotor and motor speed. Accuracy and speed for each test were *z*-transformed and summarized into an efficiency score. A factor analysis was used to summarize these efficiency scores into four factors (3), including executive function, complex reasoning, memory, and social cognition. Here, we focused on the executive function factor score. Of the sample of 946 participants with complete imaging data that passed quality assurance, two participants had incomplete cognitive data. Accordingly, 944 participants were used in the analysis examining the association between cognition and control energy.

#### Image acquisition

As previously described (4), all MRI scans were acquired on the same 3T Siemens Tim Trio whole-body scanner and 32-channel head coil at the Hospital of the University of Pennsylvania.

##### Structural MRI

Prior to dMRI acquisitions, a 5-minute magnetization-prepared, rapid acquisition gradient-echo T1-weighted (MPRAGE) image (TR = 1810ms; TE = 3.51ms; FOV = 180×240mm^2^, matrix = 192×256, effective voxel resolution = 0.9×0.9×1 mm^3^) was acquired. This high-resolution structural image was used for tissue segmentation and parcellating gray matter into anatomically defined regions in native space.

##### Diffusion MRI

dMRI scans were acquired using a twice-refocused spin-echo (TRSE) single-shot echo-planar imaging (EPI) sequence (TR = 8100ms; TE = 82ms; FOV = 240mm^2^/240mm^2^; Matrix = RL:128, AP:128; Slices: 70, in-plane resolution (x and y) 1.875 mm^2^; slice thickness = 2mm, gap = 0; flip angle = 90/180/180; volumes = 71; GRAPPA factor = 3; bandwidth = 2170Hz/pixel; PE direction = AP). This sequence used a four-lobed diffusion encoding gradient scheme combined with a 90-180-180 spin-echo sequence designed to minimize eddy-current artifacts. For dMRI acquisition, a 64-direction set was divided into two independent 32-direction imaging runs in order to ensure that the scan duration was more tolerable for young subjects. Each 32-direction sub-set was chosen to be maximally independent such that they separately sampled the surface of a sphere (5). The complete sequence consisted of 64 diffusion-weighted directions with b=1000s/mm^2^ and 7 interspersed scans where b=0 s/mm^2^. The total duration of dMRI scans was approximately 11 min. The imaging volume was prescribed in axial orientation covering the entire cerebrum with the topmost slice just superior to the apex of the brain (4).

##### Field map

In addition, a B0 field map was derived for application of distortion correction procedures, using following the double-echo, gradient-recalled echo (GRE) sequence: TR = 1000ms; TE1 = 2.69ms; TE2 = 5.27ms; 44 slices; slice thickness/gap = 4/0 mm; FOV = 240 mm; effective voxel resolution = 3.8×3.8×4 mm.

##### Scanning procedure

Before scanning, to acclimate subjects to the MRI environment, a mock scanning session where subjects practiced the task was conducted using a decommissioned MRI scanner and head coil. Mock scanning was accompanied by acoustic recordings of the noise produced by gradient coils for each scanning pulse sequence. During these sessions, feedback regarding head movement was provided using the MoTrack motion tracking system (Psychology Software Tools). Motion feedback was given only during the mock scanning session. To further minimize motion, before data acquisition, subjects’ heads were stabilized in the head coil using one foam pad over each ear and a third over the top of the head.

#### Image processing

##### Structural image processing and network node definition

The structural image was processed using FreeSurfer (version 5.3) (6), and cortical and subcortical gray matter was parcellated in native structural space according to the Lausanne atlas (7), which includes whole-brain sub-divisions of the Desikan-Killany anatomical atlas (8) at multiple spatial scales. The acquired 233-region gray matter parcellation of each subject was dilated by 2mm and then masked by the boundary of each subject’s white matter segmentation (9). Once defined for each subject, the structural parcellation atlas was co-registered to the first b=0 volume of each subject’s diffusion image using boundary-based registration (10). These parcels were then used as nodes for brain network construction. The left lateral occipital parcel was missing in 18 subjects and therefore was removed from analyses, yielding 232 brain regions that were present in all participants.

##### Diffusion image pre-processing

FSL was used for diffusion data processing (11, 12). The two consecutive 32-direction acquisitions were merged into a single 64-direction time series. In-scanner head motion and the total network strength were used as covariates in this study. Specifically, in-scanner head motion was measured by the mean relative volume-to-volume displacement between the higher SNR b=0 images (n=7), which summarizes the total translation and rotation in 3-dimensional Euclidean space (1, 9). A mask in subject diffusion space was defined by registering a binary mask of a standard fractional anisotropy (FA) map (FMRIB58 FA) to each subject’s dMRI reference image (mean *b*=0) using FSL *FLIRT*. This mask was provided as input to FSL *eddy* in addition to the non-brain extracted dMRI image. Eddy currents and subject motion were estimated and corrected using the FSL *eddy* tool (version 5.0.5: Andersson and Sotiropoulos (13)). This procedure uses a Gaussian Process to simultaneously model the effects of eddy currents and head motion on diffusion-weighted volumes, resampling the data only once. Diffusion gradient vectors were also rotated to adjust for subject motion estimated by eddy (14). After the field map was estimated, distortion correction was then applied to dMRI images using FSL’s *FUGUE* utility.

##### Probabilistic tractography and network construction

We first fitted a ball-and-sticks diffusion model for each subject’s dMRI data with FSL *bedpostx*, which uses Markov chain Monte Carlo sampling to build distributions on principal fiber orientation and diffusion parameters at each voxel (15). In contrast to tensor-based approaches, this method allowed us to model up to two crossing fibers per voxel, enhancing sensitivity to more complex white matter architecture. Probabilistic tractography was run using FSL *probtrackx*, which repetitively samples voxel-wise fiber orientation distributions to model the spatial trajectory and strength of anatomical connectivity between specified seed and target regions (15).

Each cortical and subcortical region defined along the gray-white boundary was selected as a seed region, and its connectivity strength to each of the other 231 regions was calculated using probabilistic tractography. At each seed voxel, 1,000 samples were initiated. We used the default tracking parameters (a step-length of 0.5 mm, 2,000 steps maximum, curvature threshold of 0.02). To increase the biological plausibility of white matter pathways reconstructed with probabilistic tractography, streamlines were terminated if they traveled through the pial surface, and discarded if they traversed cerebro-spinal fluid (CSF) in ventricles or re-entered the seed region (9). The connection probability from the seed voxel *i* to another voxel *j* was defined by the number of fibers passing through voxel *j* divided by the total number of fibers that were not rejected by exclusion criteria sampled from voxel *i*. For a seed cortical region, 1,000×*n* fibers were sampled (1,000 fibers per voxel), where n is the number of voxels in this region. The number of fibers passing through a given region divided by 1,000×*n* is calculated as the connectivity probability from the seed region to this given region. Therefore, a 232*232 connection probability matrix was created for each subject. Notably, the probability from region *i* to region *j* is not necessarily equivalent to the one from region *j* to region *i* due to the dependence of tractography on the seeding location. Thus, we defined the unidirectional connectivity probability *P_ij_* between region *i* and region *j* by averaging these two probabilities (9, 16).

#### Defining *a priori* network modules

Each of the 232 nodes in our network was assigned to a standard set of 7 functional systems originally defined by Yeo, *et al*. (17) in a whole-brain clustering analysis. To make this assignment, we calculated the purity index for the 7-system parcellation and brain regions from the Lausanne 232 parcellation atlas as in prior work (18). This measure quantifies the maximum overlap of cortical Lausanne labels and functional systems defined by Yeo, *et al*. (17). Each cortical Lausanne label was assigned to a functional system by calculating the non-zero mode of all voxels in each brain region (**Fig. S2**). Subcortical regions were assigned to an eighth, subcortical module.

#### Control analysis

We investigated how a structural brain network composed of white matter fiber tracts constrains the brain in transitioning from a baseline state (i.e., 1×232 zero vector) to a frontoparietal activation state, which was defined as regions in the frontoparietal system that had activity magnitude equal to 1 while other regions had activity magnitude equal to 0. According to previous studies (19–22), we employed a simplified noise-free linear continuous-time and time-invariant network model:

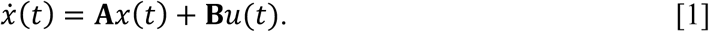

Here, *x(t)* is a 1×N vector that represents the brain state at a given time, where N is the number of ROIs (N = 232). The initial sate *x(0)* is a 1×232 zero vector, and the target state *x_T_* is a 1×232 vector of frontoparietal activation. The matrix **A** encodes the connection probability weighted network, where **A** has been scaled by its largest eigenvalue and had the identity matrix subtracted to assure that it is stable (19, 20, 22). The matrix **B** is a N×N input matrix that identifies the nodes in the control set. Here, **B** is an identity matrix because all 232 regions in the whole brain were control nodes. The input *u(t)* denotes the control energy injected for each node at a given time.

We were interested in a control task where the system transitions from initial state *x(0)* to target state *x_T_* with minimum-energy input, which is an optimal control problem. We first defined a cost function as the weighted sum of the energy cost of the transition and the integrated squared distance between the transition states and the target state.

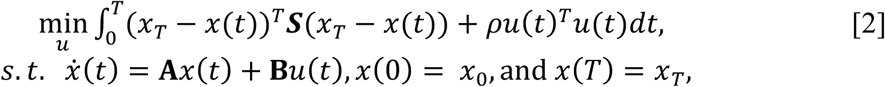

where *x_T_* is the target state (*x_T_* – *x*(*t*))^*T*^ (*x_T_* – *x*(*t*)) is the distance between the state at time *t* and the target state *x_T_*, T is a free parameter that defines the finite amount of time given to reach the target state, and *ρ* is a free parameter that weights the energy constraint. Because the time of each step was defined as 0.001, there were 1,000 steps from initial to target state if we set T=1. **S** is 0-1 diagonal matrix of size N×N that selects only the nodes that we wish to control. Here, we only constrain the activity of the frontoparietal system. Specially, (*x_T_* – *x*(*t*))^*T*^ **S**(*x_T_* – *x*(*t*)) constrains the trajectories of all nodes by preventing the system from traveling too far from the target state, and (*u*(*t*)^*T*^ *u*(*t*) constrains the amount of energy used to reach the target state.

To compute an optimal *u\** that induces a transition from the initial state *x(0)* to the target state *x_T_*, we define a Hamiltonian as:

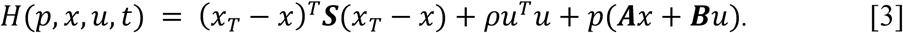

From the Pontryagin minimum principle (23), if *u\** is a solution to the minimization problem with corresponding trajectory *x\**, then there exists *p\** such that:

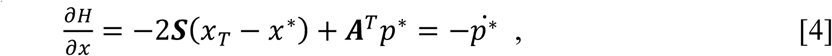

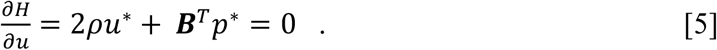

From [5] and [1], we derive that

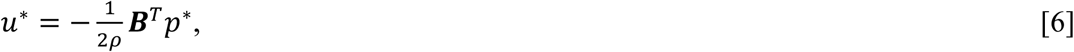

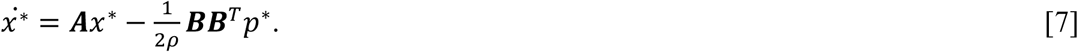

Then, we rewrite equations [4] and [7] as

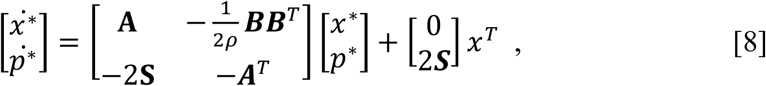

We denote:

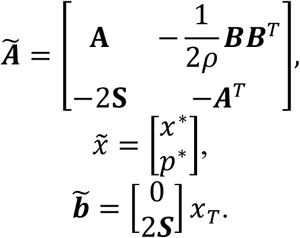

Then, equation [8] can be reduced as:

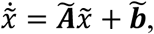

Which can be solved as:

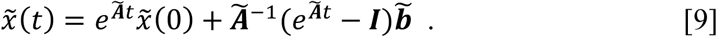

Then, by fixing t=T, we rewrote [9] as

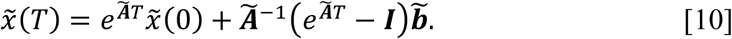

Let

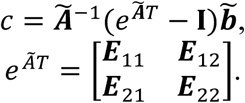

We can then rewrite [10] as:

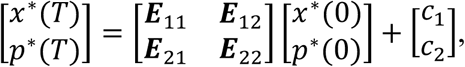

from which we can obtain

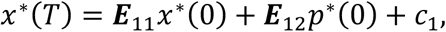

which can be rearranged to

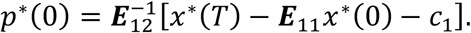

Now that we have obtained *p*(0)*, we cab use it and *x(0)* via forward to solve for 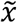 via forward integration according to Equation [9]. To solve for *u\**, we take *p\** from our solution of 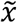 and plug it into Equation [6].

To quantify differences in trajectories, and the ease of controlling the system, we calculated a single measure of energy for every trajectory. Particularly, the energy of each control node *i* was defined as:

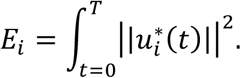

#### Prediction of brain maturity from the pattern of control energy

##### Ridge regression

A linear regression model was adopted to predict brain maturity using the pattern of whole-brain control energy. The linear model can be formalized as follows:

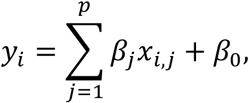

where *y_i_* is the age of the *i^th^* individual, *p* is the number of features, *x_i,j_* is the value of the *j^th^* feature of the *i^th^* subject, and *β_j_* is the regression coefficient.

To avoid over-fitting and to improve the prediction accuracy, we applied ridge regression (24–26), which used an L2 penalty during model fitting. The objective function is:

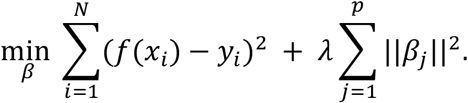

This technique shrinks the regression coefficients, resulting in better generalizability for predicting unseen samples. In this algorithm, a regularization parameter *λ* is used to control the trade-off between the prediction error of the training data and L2-norm regularization, i.e., a trade-off of penalties between the training error and model complexity. A large *λ* corresponds to a greater penalty on model complexity, and a small *λ* represents a greater penalty on training error. Compared with the traditional ordinary least squares regression, ridge regression is less impacted by multicollinearity and can avoid over-fitting (26).

##### Prediction framework

See **Fig. S6** for the schematic overview of the prediction framework. Specifically, we applied a nested 2-fold cross validation (2F-CV), with outer 2F-CV estimating the generalizability of the model and the inner 2F-CV determining the optimal parameter *λ* for the ridge regression model.

###### Outer 2F-CV

In the outer 2F-CV, all subjects were divided into 2 subsets. Specifically, we sorted the subjects according to their age and then assigned the individuals with an odd rank to subset 1 and the individuals with an even rank to subset 2 (26, 27). We first used subset 1 as a training set, and we used subset 2 as a testing set. We regressed out all covariates (i.e., sex, handedness, total brain volume, total network strength, and in-scanner head motion) from each brain feature (i.e., control energy of one brain region) for the training set, and the resulting coefficients were used to regress out the covariates for subjects in the testing set. Each feature was linearly scaled between zero and one across the training dataset, and the scaling parameters were also applied to scale the testing dataset (26, 27). We applied an inner 2-fold cross validation (2F-CV) within training set to select the optimal *λ* parameter. Based on the optimal *λ*, we trained a model using all subjects in the training set, and then used that model to predict the age of all subjects in the testing set. Analogously, we used subset 2 as a training set and subset 1 as a testing set, and repeated the above procedure. Across the testing subjects for each fold, the correlation and mean absolute error (MAE) between the predicted and actual age was used to quantify the prediction accuracy. Here, we used the scikit-learn library to implement ridge regression (http://scikit-learn.org) (28).

###### Inner 2F-CV

Within each loop of the outer 2F-CV, we applied inner 2F-CVs to determine the optimal *λ*. Specially, the training set for each loop of the outer 2F-CV was further partitioned into 2 subsets according to their rank of the age, as like the outer loop (i.e., subjects with odd rank in subset 1 and subjects with even rank in subset 2). One subset was selected to train the model under a given *λ* in the range [2^−10^, 2^−9^, …, 2^4^, 2^5^] (i.e., 16 values in total) (26), and the remaining subset was used to test the model. This procedure was repeated 2 times such that each subset was used once as the testing dataset, resulting in 2 inner 2F-CV loops in total. For each inner 2F-CV loop, the correlation *r* between the actual and predicted age and the mean absolute error (MAE) were calculated for each *λ*, and averaged over each fold. The sum of the mean correlation *r* and reciprocal of the mean MAE was defined as the inner prediction accuracy, and the *λ* with the highest inner prediction accuracy was chosen as the optimal *λ* (26, 27). Of note, the mean correlation *r* and the reciprocal of the mean MAE cannot be summed directly, because the scales of the raw values of these two measures are quite different. Therefore, we normalized the mean correlation *r* and the reciprocal of the mean MAE across all values and then summed the resultant normalized values.

###### Randomly split 2F-CV

In the above prediction analysis, we split subjects into two halves according to their age rank. For completeness, we also split the subjects randomly into two halves for both outer 2F-CV and inner 2F-CV, and calculated the mean correlation *r* and MAE across two folds. Because the split is random, we repeated this procedure 100 times and averaged the correlation and MAE across the 100 times to acquire the final prediction accuracy.

#### Specificity and sensitivity analysis

##### Controlling for modal controllability

The present work explored a specific transition of the brain from a baseline state to a state of frontoparietal activation state by enacting multi-point control. In contrast, modal controllability quantifies the difficulties of transitioning to all possible states via single-node control (29). Modal controllability identifies brain areas that can push the brain into difficult-to-reach states; our prior work has shown that modal control increases with age in youth (30). Accordingly, it is important to establish whether our present results were driven by developmental changes in modal controllability. As in Tang, *et al*. (30), before calculating controllability, we scaled the matrix by 1+ξ_0_, where ξ_0_ is the largest eigenvalue value of the matrix. Next, we conducted sensitivity analyses where we controlled for modal controllability by including it as a covariate in the regression equation at each resolution of analysis (e.g., whole brain, functional system, network nodes).

##### Controlling for overall network modularity

Given that structural brain networks are modular, and modularity changes with age (31), it is important to evaluate if observed developmental associations with control energy might be driven by changes in network modularity. We calculated network modularity quality (*Q*) using the community structure defined by the functional atlas (17). See Baum, *et al*. (18) for the details of the calculation of *Q_Yeo_*, where we have shown that *Q_Yeo_* was highly similar to the network modules identified using data-driven community detection procedures (32, 33). We scaled the matrix by the maximum eigenvalue before calculating *Q*, as the matrix was scaled when calculating the energy. We do not subtract the identity matrix because it would lead to (uninterpretable) negative values of *Q*. Finally, we controlled for *Q* by including it as a model covariate in sensitivity analyses, which were conducted at all resolutions (whole brain, functional systems, and network nodes).

**Fig. S1.**
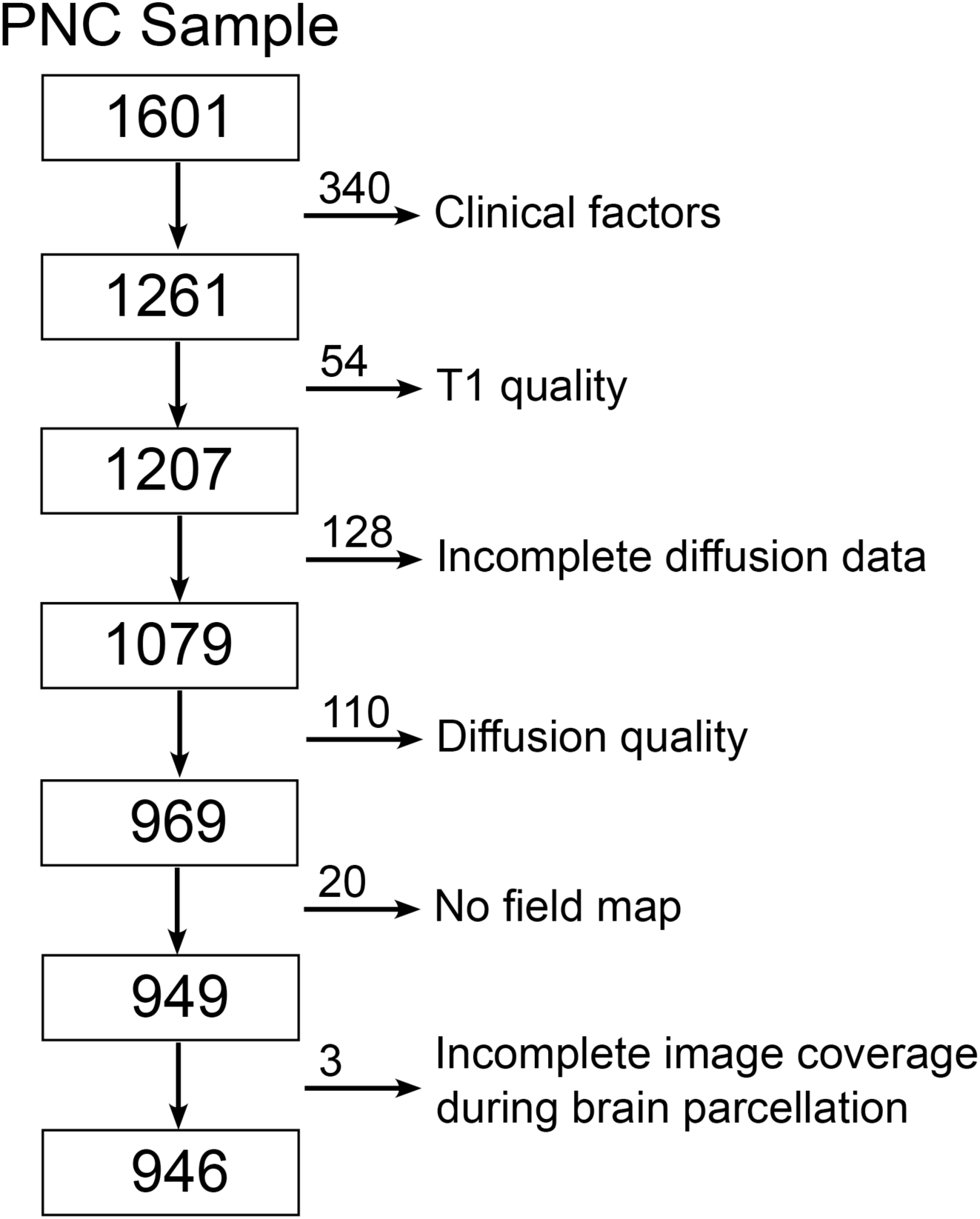
Sample construction. The cross-sectional sample of the Philadelphia Neurodevelopmental Cohort (PNC) has 1601 participants in total. 340 subjects were excluded owing to clinical factors, such as medical disorders. Then, 312 subjects were excluded because of low quality of T1 or diffusion data, incomplete diffusion data, lacking of field map. Finally, 3 subjects were excluded due to incomplete image coverage during brain parcellation. The final sample consisted of the remaining 946 subjects.

**Fig. S2.**
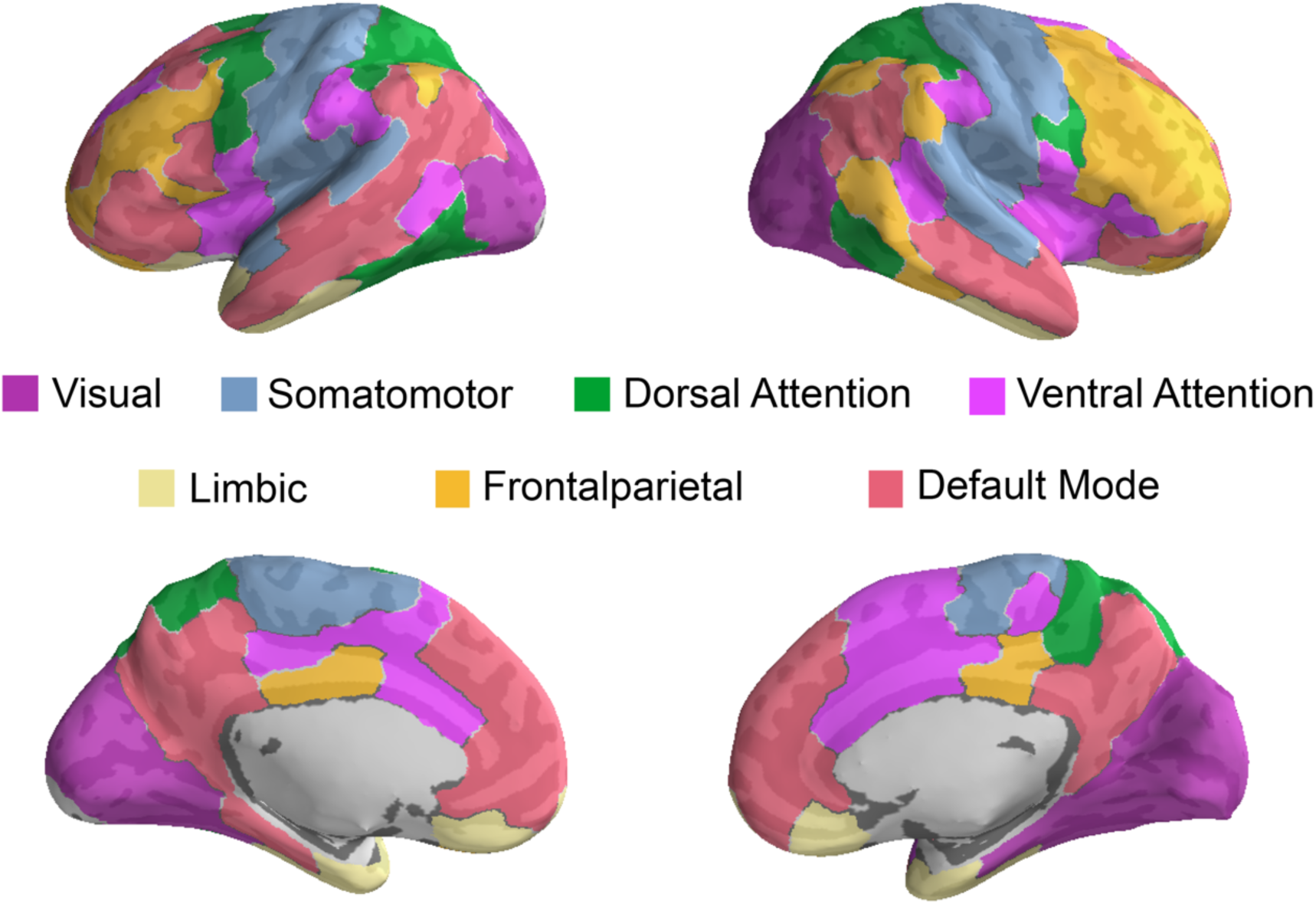
Functional brain networks defined by (17). Each parcel was mapped to one of these networks.

**Fig. S3.**
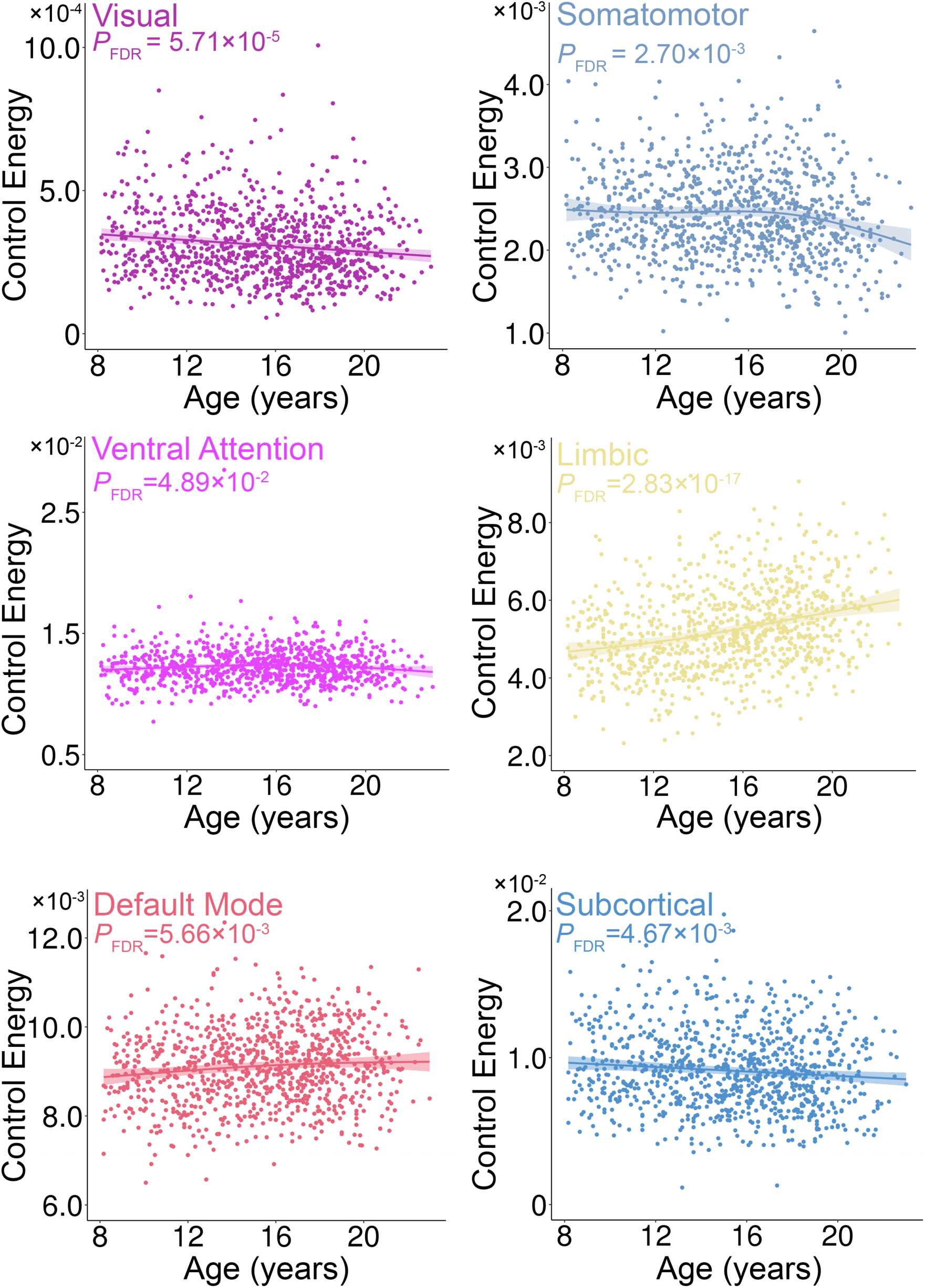
Scatter plots of significant age effects of control energy at the system scale. The control energy of visual, somatomotor, and subcortical systems decline significantly with age, while that of ventral attention, limbic and default mode systems increase significantly with age. Data points represent each subject (n = 946), the bold line indicates the best fit from a general additive model, and the shaded envelope denotes the 95% confidence interval.

**Fig. S4.**
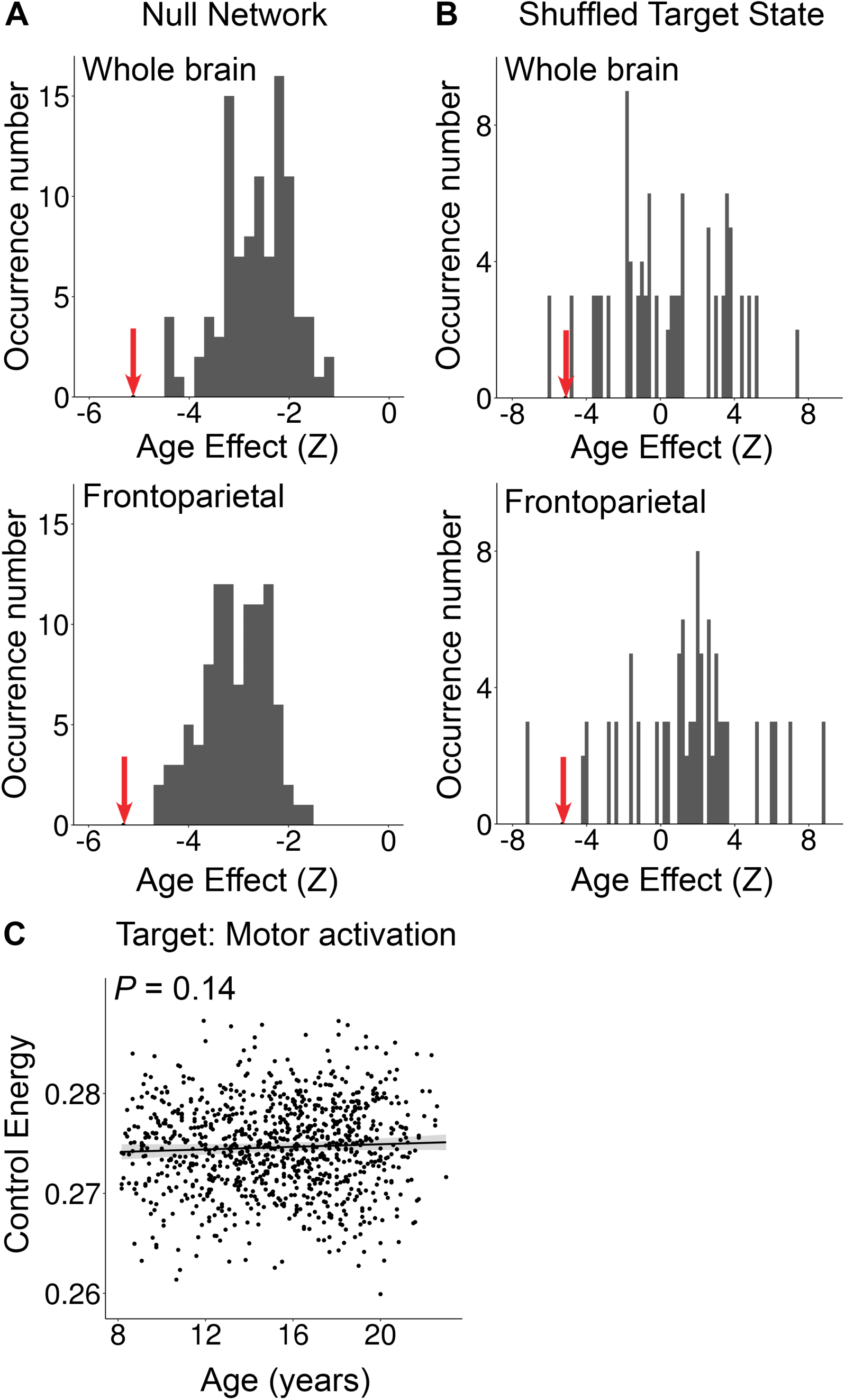
Specificity and sensitivity analyses provide convergent results. **A)** The distribution of the age effect of average control energy of whole-brain and frontoparietal systems when using null model networks, which preserve the degree and strength distribution. The red arrow indicates the actual age effect estimated using the data from the real brain network. **B)** The distribution of the age effect of control energy when the activation target states were shuffled uniformly at random. The red arrow indicates the actual age effect with the real target state. **C**) The control energy required to reach a motor activation state did not change over the age range studied.

**Fig. S5.**
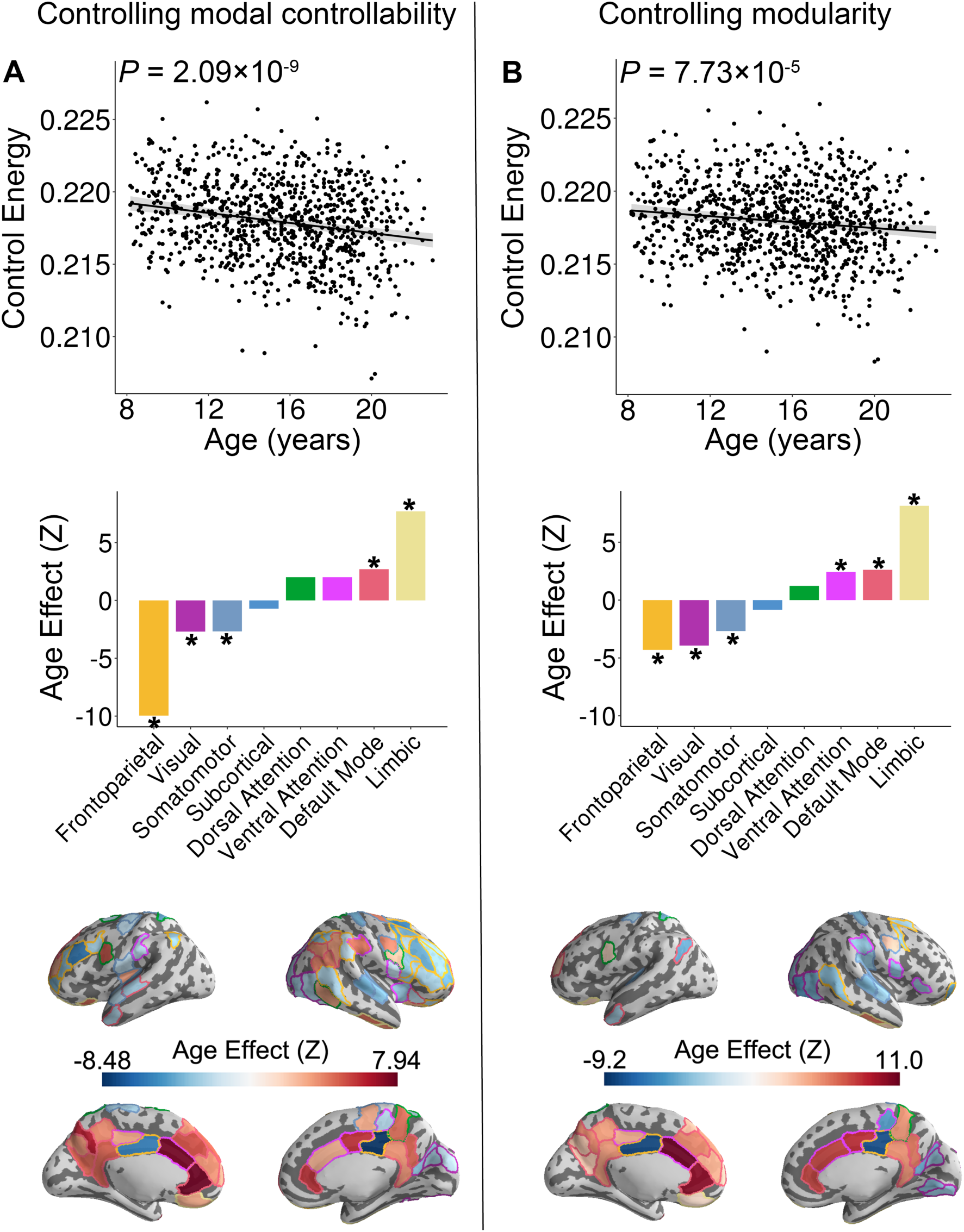
Age effects at the whole-brain, cognitive system, and nodal level are maintained after controlling for the **A)** modal controllability and for the **B)** network modularity.

**Fig. S6.**
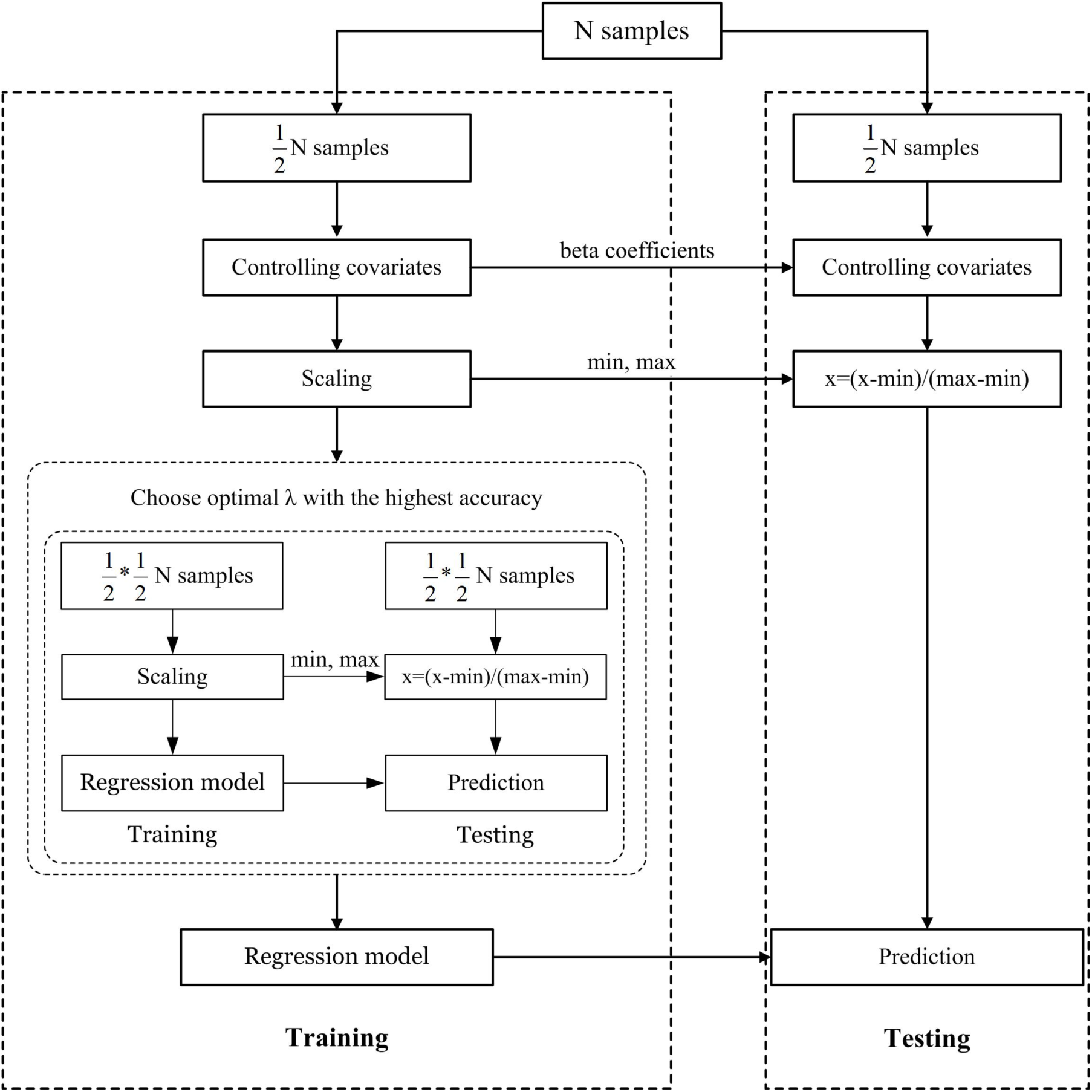
Schematic overview of one outer loop of the nested 2-fold cross-validation (2F-CV) prediction framework. All subjects were divided into 2 halves according to age rank, with the first half used as a training set and the second half used as a testing set. We regressed out covariates from each brain feature in the training set, and the resulting coefficients were used to regress out the covariates for subjects in the testing set. Each feature was linearly scaled between zero and one across the training dataset, and the scaling parameters were also applied to scale the testing dataset. An inner 2F-CV was applied within training set to select the optimal *λ* parameter. Based on the optimal *λ*, we trained a model using all subjects in the training set, and then used that model to predict the age of all subjects in the testing set.

**Fig. S7.**
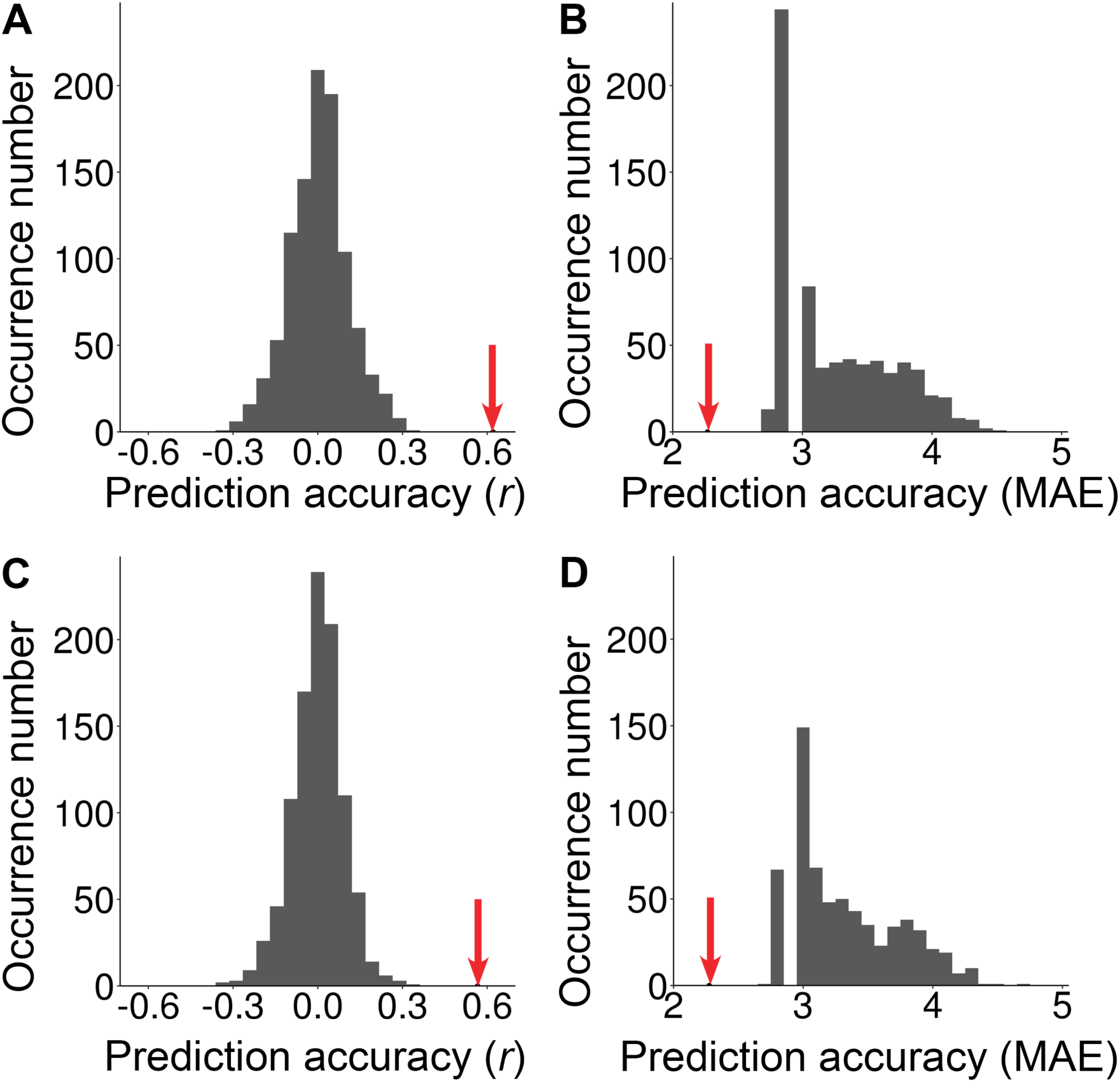
The histograms of the permutation distribution of the **A)** correlation *r* and **B)** MAE with the first subset used as a training set and the second subset used as a testing set, and **C)** correlation *r* and **D)** MAE with the first subset used as the testing set and the second subset used as training set. The red arrow represents the actual prediction accuracy (i.e., *r* or MAE). The actual correlation *r* was significantly higher than expected by chance (p<0.001) and the actual MAE was significantly lower than expected by chance (p<0.001).

